# High temporal resolution of gene expression dynamics in developing mouse embryonic stem cells

**DOI:** 10.1101/084442

**Authors:** Brian S. Gloss, Bethany Signal, Seth W. Cheetham, Franziska Gruhl, Dominik Kaczorowski, Andrew C. Perkins, Marcel E. Dinger

**Affiliations:** Garvan Institute of Medical Research, Sydney, Australia; St Vincents Clinical School, Faculty of Medicine, UNSW Australia, Sydney, Australia; The Gurdon Institute and Department of Physiology, Development, and Neuroscience, University of Cambridge, Cambridge, United Kingdom; Center for Integrative Genomics, University of Lausanne, Lausanne, Switzerland; Mater–UQ Research Institute, The University of Queensland, Translational Research Institute, Brisbane, Australia

**Keywords:** Bioinformatics, systems biology, lncRNAs, embryonic stem cell development, genome biology

## Abstract

Investigations of transcriptional responses during developmental transitions typically use time courses with intervals that are not commensurate with the timescales of known biological processes. Moreover, such experiments typically focus on protein-coding transcripts, ignoring the important impact of long noncoding RNAs. We evaluated coding and noncoding expression dynamics at high temporal resolution (6-hourly) in differentiating mouse embryonic stem cells and report the effects of increased temporal resolution on the characterization of the underlying molecular processes. We present a refined resolution of global transcriptional alterations, including regulatory network interactions, coding and noncoding gene expression changes as well as alternative splicing events, many of which cannot be resolved by existing coarse developmental time-­-courses. We describe novel short lived and cycling patterns of gene expression and temporally dissect ordered gene expression at bidirectional promoters and responses to transcription factors. These findings demonstrate the importance of temporal resolution for understanding gene interactions in mammalian systems.

**Links to data:** Data has been deposited into GEO: The Reviewer access link is: http://www.ncbi.nlm.nih.gov/geo/query/acc.cgi?token=cnglummejbkltyj@acc=GSE75028

## Introduction

Over the past decade, transcriptomic investigations into the of nature embryonic stem cell (ESC) differentiation have elucidated key biochemical features of stemness and differentiation. Increasingly, it has become apparent that understanding the dynamics and coordination of gene expression signatures over time during the key phases of differentiation is critical to adequate characterization of fundamental biological processes.

ESC differentiation in mouse is a highly complex cascade of gene expression changes that allow single pluripotent cells in culture to progress to an organoid resembling a pre-implantation blastocyst within only five days. The spontaneous differentiation of these cells in culture has provided key insights into the developmental processes underlying the generation of the primary germ cell layers(1). Microarray and RNA sequencing have provided a means to characterize the molecular transitions in gene expression underlying ESC biology and more recently single cell transcriptomic studies have provided the first glimpses into the molecular history of these cells(2). However, it is clear that much more of the transcriptional landscape of ESC remains to be elucidated(3).

Access to new technologies, such as massively parallel sequencing (MPS), has led to a dramatic increase in our knowledge of the mammalian transcriptome. Early genomic tiling array analysis indicated that most of the genome was transcribed into RNA(4). MPS of the transcriptome validated this observation and revealed that the majority of the mammalian genome is pervasively transcribed as interlaced and overlapping RNAs(5), many of which lack protein-coding potential(6). The large number of long-noncoding transcripts (lncRNA) has become the focus of significant interest due to their exquisite cell type specific expression(7), potent biological function (8,9), and rapid transactivation of cellular processes. However, in general, lncRNAs are lowly expressed and short lived(10), possibly because, unlike mRNAs that require translation, are able to exert their function directly. These qualities obfuscate their identification and characterization with traditional approaches that are tuned to the properties of mRNAs (11). Owing to the relative infancy of the field, the vast majority of noncoding transcripts are of unknown function(12). Additionally, the expression patterns of these genes imply that their function is dependent on cellular context and likely regulatory(8), thus the identification of these molecules and the context in which they act remains a research priority(13).

Various expression profiling studies, using both microarrays and RNA-seq(14-17), have been used to explore the molecular changes occurring during ES cell development, typically at 24-hourly or more. This potentially has lead to incomplete gene expression relationships through the phenomenon of temporal aggregation bias whereby each time point is assumed to represent all the signaling changes occurring in that time window (18). In contrast to single cell based approaches-which provide insight into the state of individual cells - examinations of whole cell populations provides system-wide behavior and a practical means to explore gene expression dynamics across time. The combination of these techniques has recently shed light the molecular framework of cellular differentiation (19). Higher temporal resolution has also shown rapid induction (within two hours of retinoic acid stimulation) of lncRNAs associated with the HOX locus (20). Furthermore, high temporal resolution has provided valuable insights into transcriptional annotation and regulation in drosophila (21,22), Xenopus (23) and C.elegans (24).

Here we show that additional temporal resolution of the global transcriptome in spontaneously differentiating mESC cells following LIF withdrawal enables the capture of the rapid and complex dynamic regulatory and noncoding changes occurring during ES development. We analyzed the transcriptome of differentiating mouse ESCs at six-hourly intervals over a five-day period, over which time the three primordial germ layers are specified. Using this fine-resolution temporal sampling approach, we identify significant transitions in the transcriptome and large-scale shifts in observable transcription factor activities that could not be observed at 24 hourly sampling periods. Moreover, we identify entirely novel coding and noncoding transcripts that are expressed only within specific sub-24-hour window. By leveraging the high sampling frequency of the data, we are able to both accurately recapitulate known regulatory cascades in ES development and predict and refine others. Finally, using correlative approaches, we can infer functions for uncharacterized lncRNAs and predict the regulatory centers across the genome that coordinate early development.

## Materials and Methods

### Sample Generation and Library Preparation

Biological duplicate, low passage number (P18) W9.5 ESCs were cultured and differentiated as described previously(14,25). Cultures were harvested every six hours from the induction of differentiation to 120 hours post differentiation induction. Total RNA from cultures was purified using Trizol (Life Technologies) and DNase treatment was performed by RQ1 DNase (Promega) according to the manufacturer’s instructions. RNA integrity was measured on a Bioanalyzer RNA Nano chip (Agilent). RNA-Seq library preparation and sequencing of Poly-A-NGS libraries generated from 500 ng total RNA using SureSelect Strand Specific RNA Library Preparation Kit (Agilent) according to the manufacturer’s instructions. Paired-end libraries were sequenced to the first 100 bp on a HiSeq 2500 (Illumina) on High Output Mode.

### Quality control and read mapping

Library sequencing quality was determined using FastQC (Babraham Bioinformatics) and FastQ Screen (Babraham Bioinformatics). Illumina adaptor sequence and low quality read trimming (read pair removed if < 20 base pairs) was performed using Trim Galore! (Babraham Bioinformatics: www.bioinformatics.babraham.ac.uk/). Tophat2 (26) was used to align reads to the December 2011 release of the mouse reference genome (mm10) as outlined by Anders et al.(27). Read counts data corresponding to GENCODE vM2 transcript annotations were generated using HTSeq (28). de novo transcript assembly was performed on each merged BAM file using Cufflinks’ reference annotation based transcript (RABT) assembly(29), using the Gencode vM2 transcriptome(30) as a guide (options: -u -I 500000 -j 1.0 -F 0.005 –trim-3-dropoff-frac 0.05 –g gencode.vM2.annotation.gtf –library-type fr-firststrand). Transcript assemblies were then merged using Cuffmerge(31) using default parameters, and compared to the Gencode vM2 reference transcriptome using Cuffcompare(31). Novel transcripts with a Cuffcompare class code of j, i, o, u or x were filtered using three steps to find novel lncRNAs. First, a Browser Extensible Data (BED) format file was generated using a python script (https://gist.github.com/davidliwei/1155568) and any single exon transcripts were removed. Second, the FASTA-formatted sequence for each transcript was obtained using BEDTools(32), the nucleotide (nt) length and open reading frame (ORF) size found using Perl scripts, and those with a length less than 200 nt or a ORF size greater than 300 nt were removed. Lastly, transcript sequences were submitted to Coding Potential Calculator (CPC)(33), and those with a coding potential of >0 were removed.

### Bioinformatics

All analyses were performed in the R Statistical Environment(34). Briefly, counts data were background corrected and normalized for library size using edgeR(35), then transformed using voom(36) for differential expression analysis using LIMMA(37). Transcription Factor (TF) activity was inferred from gene expression data using DREM(38) with a branching P-value of 0.001 based on curated TF-target gene lists associated with mouse ESC differentiation from ChEA(39). TF-target gene was calculated by maximal Pearson’s correlation coefficient of >0.8 using a custom autocorrelation analysis and verified with the “ccf” function in R. Gene differential exon (DEX) usage was analyzed by DEXSeq(40) on vM2 gene annotations using default settings and an adjusted p value cutoff of 0.001 for DEX between biological duplicates at each consecutive time-point. Genome position analyses were performed using genomic ranges(41) based on vM2 annotations imported with ‘rtracklayer’(42) and Pearson’s correlation coefficient of gene expression Bidirectional genes were defined as two genes with expression data on opposing strands with <2000 bp between the transcriptional start sites (TSS). Co-expressed gene clusters were defined as >5 contiguous genes with expression data displaying a Pearson’s Correlation Coefficient of >0.5 with neighbouring genes. Cluster co-expression data was visualized with corrplot(43) and Cytoscape (v3.1.0(44)), location of related clusters was visualized by Circos(45). Gene expression periodicity was measured on 120 interpolated expression values(46) for each replicate time series using GeneCycle(47), candidate periodically expressed genes were identified as having the same calculated dominant cycling frequency between biological replicates. Time-dependent expression signatures were established using maSigPro(48) with a replicate correlation coefficient cutoff of 0.8. Target genes of potential regulatory (top 50 most highly and/or variably expressed) lncRNAs were identified using the GeneReg package(49) on 100 point-interpolated expression data based on fitted expression values between duplicates and setting a maximum time delay of 18 hours and a global correlation coefficient of 0.9 and visualized using Cytoscape. Gene lists were functionally annotated with KEGG and Reactome pathways (adjusted p value <0.05) using the clusterProfiler and ReactomePA packages(50).

## Results

### The dynamic transcriptome of mESC differentiation at high temporal resolution

A median 42-million, paired-end 100-bp reads (Supplementary Figure S1A) were mapped from stranded, poly-A derived cDNA libraries derived from biological duplicate, six-hourly time courses of mESC differentiation over five days where key differentiation programs occur (0-120 hours, Figure 1A). Transcript-level expression data was generated as previously described (27), then normalized for library size and transformed for data visualization and differential gene expression analysis. Evaluation of 24 hourly time points indicated that our data was comparable to previously published data in a similar model (51) (Supplementary Figure S1B). An interactive gene expression portal was created to visualise this data (https://betsig.shinyapps.io/paper_plots).

**Figure 1.**
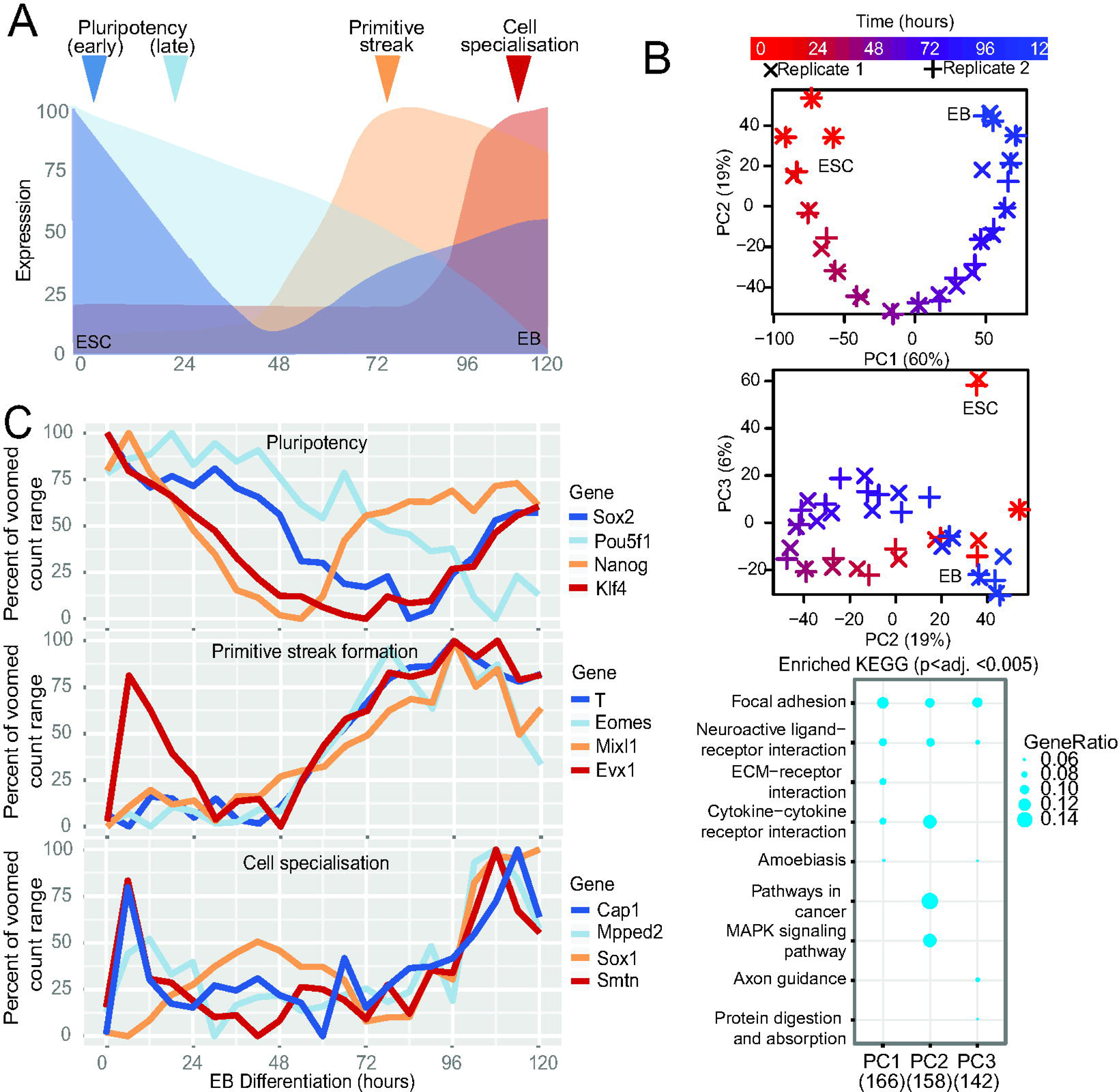
Global and gene-specific evaluation of augmented temporal resolution in mES differentiation. **(A)** Schematic of mouse embryonic stem cell (ESC) differentiation into embryoid bodies (EB) over the time course evaluated here. (**B**)Analysis of the top three principle components (PCs) based on the 2,000 most variable genes from biological duplicate-6 hourly transcriptomes.**and** KEGG pathway enrichment for 500 genes contributing most to each of the top three PCs. (**C)** Expression profiles of genes associated with pluripotency, primitive streak formation and cell specialization.

To assess the reproducibility and provide confidence in the biological validity of the global transcriptome trends, a principle components analysis (PCA) was performed on the 2,000 most variable genes (Figure 1B). This analysis indicated that biological replicates clustered closely, indicating that synchrony was retained, and that the major contributor to the determination of variance was explained by time. Deconvolution of the dimensions yielded time-dependent expression (in the first dimension) of genes enriched in focal adhesion/ ECM interactions KEGG pathways. Interestingly, the second dimension deconvolution (PC2), in which undifferentiated ESCs resemble the more differentiated embryoblast, yielded genes enriched in MAPK-signaling and cancer pathways, implying that the process of differentiation involves a partial retention of a cells capacity for self renewal. In the third component (PC3), in which the undifferentiated ES cell is separate, the axon-guidance pathway was enriched. We then evaluated expression patterns of genes associated with pluripotency, primitive streak formation and cell specialization (Figure 1C). We observed that, although the gene expression patterns were broadly consistent with published studies (Supplementary Figure S1B), there were changes in expression on less than 24 hourly timeframes that could not be attributed to measurement biases (within the top 5% of deviation from loess-smoothed expression values). To establish how prevalent sub-24 hour gene expression changes were in in the transcriptome of developing ESCs, we evaluated the extent to which gene expression patterns observed 24 hourly were unable to capture gene expression changes happening within that window (temporal aggregation bias (18)). We observed that, compared to 24 hour time points, 417 more genes had counts data considered sufficient for differential gene expression analysis; reflecting a substantial increase in detected noncoding genes over protein coding (>12% vs. 2% respectively, chi-squared p value <0.001, Supplementary Table S1, Supplementary Figure S1C). Furthermore, the additional time points allowed the assembly of 58% more novel multiexonic intergenic, antisense and intronic noncoding RNAs from the data - indicating that a substantial proportion of noncoding transcripts are present on timescales much shorter than 24 hours. Finally, to ensure that the 6-hourly measures represented distinct gene expression patterns to the 24-hourly measures, we observed that no single 24-hourly measure was representative of the average expression over that day (Mann-Whitney U p. adj. <1E-145) and that more than 1,000 genes displayed a more than 2-fold difference mostly in the first 24 hours of differentiation (Supplementary Figure 1D-E). These results indicate that enhanced temporal resolution reduces the phenomenon of temporal aggregation bias and allows the observation of more distinct cell expression states than typical time-courses.

### An improved signaling cascade described by higher temporal resolution

Increased sampling frequency can provide a powerful insight into understanding of the contribution of gene regulatory networks to cellular differentiation (21). We utilized the DREM v2 analysis tool (38) to evaluate transcription factor (TF) target gene expression patterns. Divergence of gene targets responsive to groups of TF at each time point, either 24-hourly or 6-hourly (Figure 2A-B) was shown if the overall difference was significant at p<0.001. Compared to 24-hourly, the observed complexity was significantly higher, especially in the first 48 hours. We observed that significant changes in gene regulation occurred continuously within the 24-hour windows. Most notably, first 24 hours following depart from pluripotency resembles an ordered cascade of TF activity (Figure 2A, Supplementary Figure S2A) with large-scale changes in TF activity at 12, 18 and 24 hours; of which little can be deduced measuring at just 24 hours (Figure 2B, Supplementary Figure S2B). Focusing on the interplay between two key transcription factors (Otx2 and Pou5f1/Oct4(52), Figure 2A), we observed a rapid rise in Otx2 activity in the first six hours and stable Pou5f1activity for the first 24 hours (Red Box). Otx2 activity did not coincide with mRNA expression of the factor itself (Figure 2C), although previous studies have observed increased in Otx2 protein expression within 3-hours of differentiation(52), however periodic drops in Pou5f1 mRNA expression appeared to coincide with decreases in POU5F1 target genes, we calculated the time taken for Pou5f1 expression to result in changes in highly positively correlated (r>0.8) target genes using a cross-correlation approach similar to (53). We then evaluated how these “delays” enriched for certain Reactome pathways (Figure 2D). We found rapid effects for targets enriched for “gene expression”- and a delayed effect on “cell cycle” pathways compared to a null distribution produced by 500 random “target” selections (grey). These were similarly observed in the DREM GO-term enrichment tool for Pou5F1 targets decreasing in expression at 42 (early-Transcription Factor Activity) and 54 hours (late-Epithelial Proliferation; Figure 2A, Blue Box & Supplementary Figure S2C) and associated with the decrease in Pou5F1 expression (Figure 2C, Blue Box). Importantly, Pou5F1 mRNA and protein expression are temporally correlated (52). This result implies that TF-target genes may be activated in an ordered-time dependent fashion. To explore this more broadly, we evaluated other TF-target gene temporal dynamics for other TFs that exhibited strong positive or negative correlations between the TF and their target genes. We found evidence of highly structured TF-target expression patterns in time for negatively correlated Pou5f1 and Suz12 targets, as well as positively correlated Nanog, Myc, Sox2 and Suz12 targets (Supplementary Figure S3).

**Figure 2.**
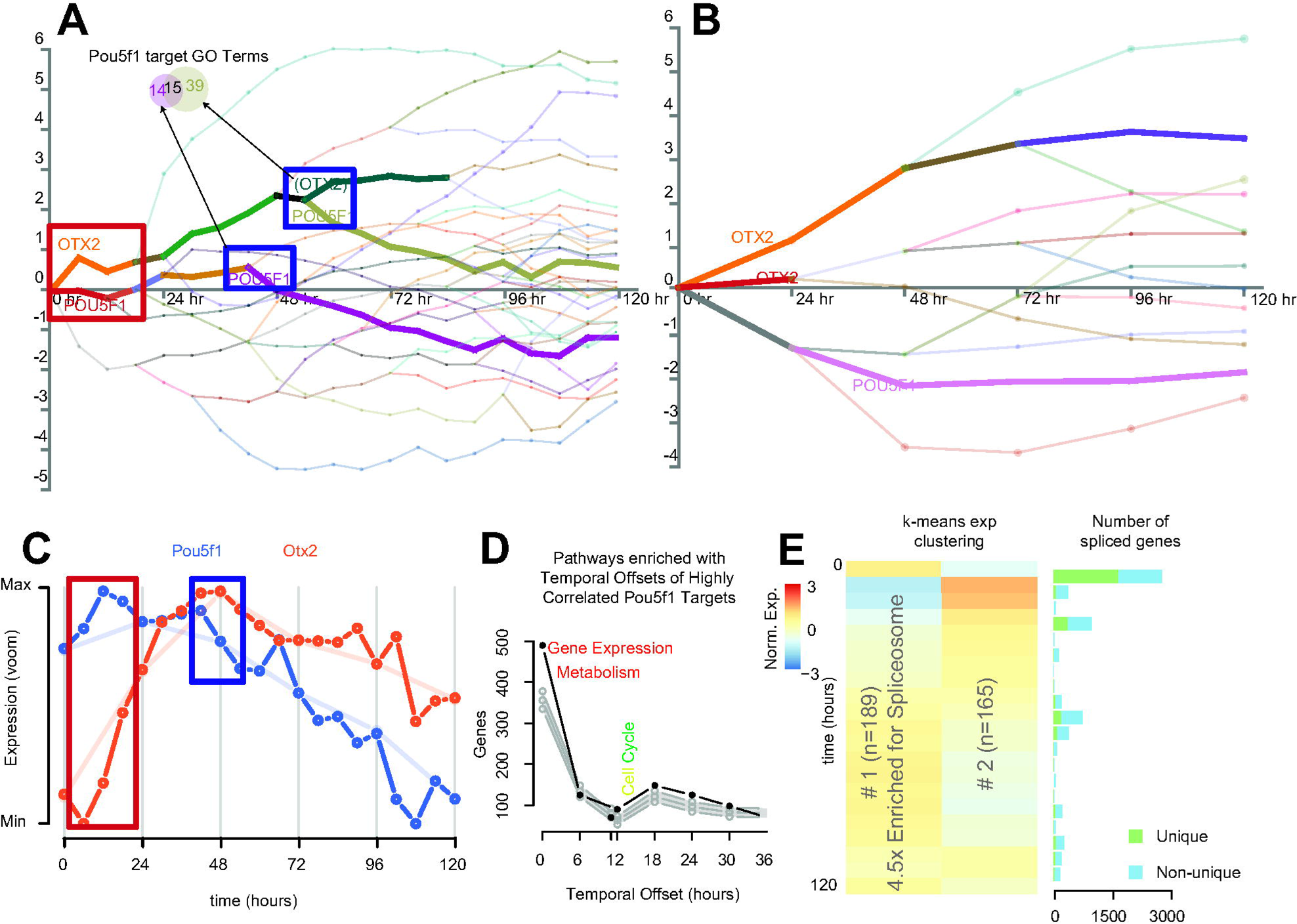
Insights into regulatory and gene expression kinetics. **(A & B)** Observable regulatory network dynamics at 24- and 6-hourly measures with OTX2 and POU5F1 target containing profiles annotated and in bold, See Supplementary Figure S2 for full figure. Transcriptomes at 24- (top) and 6-hourly (bottom) were subjected to DREM analysis of mouse TF/target gene interactions. A p-value cutoff of 0.001 was applied to calculating divergent TF activity (splits). Relative circle sizes are proportional to the spread of gene expression levels corresponding to that point. Red and blue boxes pertain to branch points of interest (**C)** Expression of the key transcription factors Pou5F1 (Oct4) and OTX2. Red and blue boxes correspond to the time points highlighted in part A. (**D)** Distribution of the number of genes and the time delay required to meet a maximum correlation (>0.8) between gene targets of Pou5f1 and pPou5f1 itself compared to 95% quantiles of 500 random gene selections. (**E)** Two k-means clusters short-lived RNA (slRNA) genes displaying differential expression without changes at 24-hourly time points (adj. p<0.0001).

These observations of precise temporal ordering of transcriptional events emphasize the importance of factoring time delays into understanding gene regulatory networks (54) and highlight the capacity of increased temporal resolution to directly identify –rather than inference in most cross-correlation approaches-valuable new knowledge of regulator-target gene interactions.

### Increased temporal resolutions identifies genes with previously uncharacterized expression patterns (Short-lived (slRNA) & Cycling (cycRNA))

Having established that the increased temporal resolution markedly improves the molecular framework for evaluating the contribution of gene expression to ES differentiation, we next sought to identify gene expression signatures previously unable to be resolved using lower temporal resolution. For each 24-hour period, we identified genes that were differentially expressed between 0 and 6, 12 and 18 hours but not between any 24-hourly measures (Supplementary Figure S2D). We identified 1,135 genes with significant changes in gene expression that were unchanged between any 24-hourly comparison (adjusted p<0.0001). Of these, 354 were differentially expressed for more than half of the corresponding 24-hour window, mostly in the first and last 24-hour periods. These genes were described as short-lived RNAs (slRNAs). slRNA expression patterns over the first 24 hours of differentiation were found to be positively correlated with the same time window of retinoic acid directed differentiation (20) (Supplementary Figure S2E) implying that these genes may form part of the early response to differentiation signals. K-means clustering and KEGG pathway analysis of the expression profiles of these genes (Figure 2E) revealed enrichment in genes associated with the spliceosome (p=0.02) dramatically decreasing in expression over the first 24 hours before returning slowly to baseline. To examine whether this impacted gene-splicing patterns, we employed a differential exon (DEX) analysis between consecutive six-hourly time points and counted the number of genes displaying DEX usage (Figure 2E). Consistent with previous studies, the alternate splicing was most highly associated with cell differentiation(55) (Figure 2E). Increased temporal resolution has elucidated that these changes happen very rapidly (majority of changes in the first six hours), and that slRNAs may be involved in suppressing the alternate splicing of genes and limiting transcriptional plasticity.

Some slRNAs appeared to have periodic expression profiles. We thus sought to uncover periodic expression patterns genome-wide, by applying a fast-Fourier transformation to our data (see Methods). Periodogram analysis was utilized to ascertain the dominant cycling period for each gene. We found 137 genes, which we termed cycling RNAs (cycRNAs), sharing the same dominant cycling period of less than 36 hours in both biological replicate experiments (Supplementary Table S2). Supporting the efficacy of the approach, we found Clock, which encodes a key regulator of circadian rhythm in mammals, to have a period of 24.2 hours. We identified 20 genes that displayed characteristics of both slRNAs and cycRNAs (Supplementary Figure S2F), including Ewsr1 and Clk1, involved in gene splicing(56,57) as well as five uncharacterized lncRNAs. Given the highly specific expression patterns in this context, we propose these genes may similarly have roles in maintaining or establishing biological rhythms. Together these investigations show that the augmented temporal resolution approach provides access to gain insights from regulatory pathways by identifying transitions in expression that would otherwise have remained hidden.

### Increased temporal resolution gives insight into local gene regulation in the genome

Evaluating gene transcription at high temporal resolution in a highly dynamic process such as ES development, we anticipated that it might be feasible to dissect structural gene regulation within a given locus. To explore this possibility, we examined expression arising from transcripts that are oriented head-to-head as so-called bidirectional pairs (58,59). Interestingly, we observed that the antisense transcript for Evx1 (Figure 1C) displayed a previously unobserved (14) increase in expression in the first 24 hours after departure from pluripotency that was reflected in its paired protein coding gene Evx1 (Supplementary Figure S4A), highlighting the increased power of frequent sampling over time. In total, we identified 1,251 gene pairs with bidirectional transcriptional start sites (TSS) within 2,000 bp and evaluated correlation coefficients across the time course, distance between TSS and median expression values. Consistent with other studies, we found expression correlation more positive for bidirectional gene pairs than random transcript pairs(58) (Supplementary Figure S4B). We were also able to show that the distance between TSS of highly correlated bidirectional gene promoters is typically less than 500 bp (Figure 3A), consistent with a common regulatory domain. Highly correlated or anti-correlated genes pairs displayed differences in total gene expression, particularly with discordant gene biotypes (Figure 3B). We found that protein coding gene pairs were more likely to be of similar expression levels and positively correlated (p<0.05) than protein coding/noncoding pairs (Supplementary Figure S4C). Applying a variant of the temporal offset analysis used to measure TF-gene target delays, we calculated the time taken and defined the apparent driver gene type for peak correlation in coding/noncoding bidirectional pairs (Supplementary Figure S4D, E). This did not reveal a generalized bias in either time taken or particular “driving” gene type. However, this approach shows that the lncRNA Hotairm1, required for activation of Hoxa1(60), appears to have a six-hour delay between its expression changes and HoxA1. We present evidence of other examples of lncRNA-led expression of protein coding genes in small numbers of bidirectional pairs (Supplementary Figure S5).

**Figure 3.**
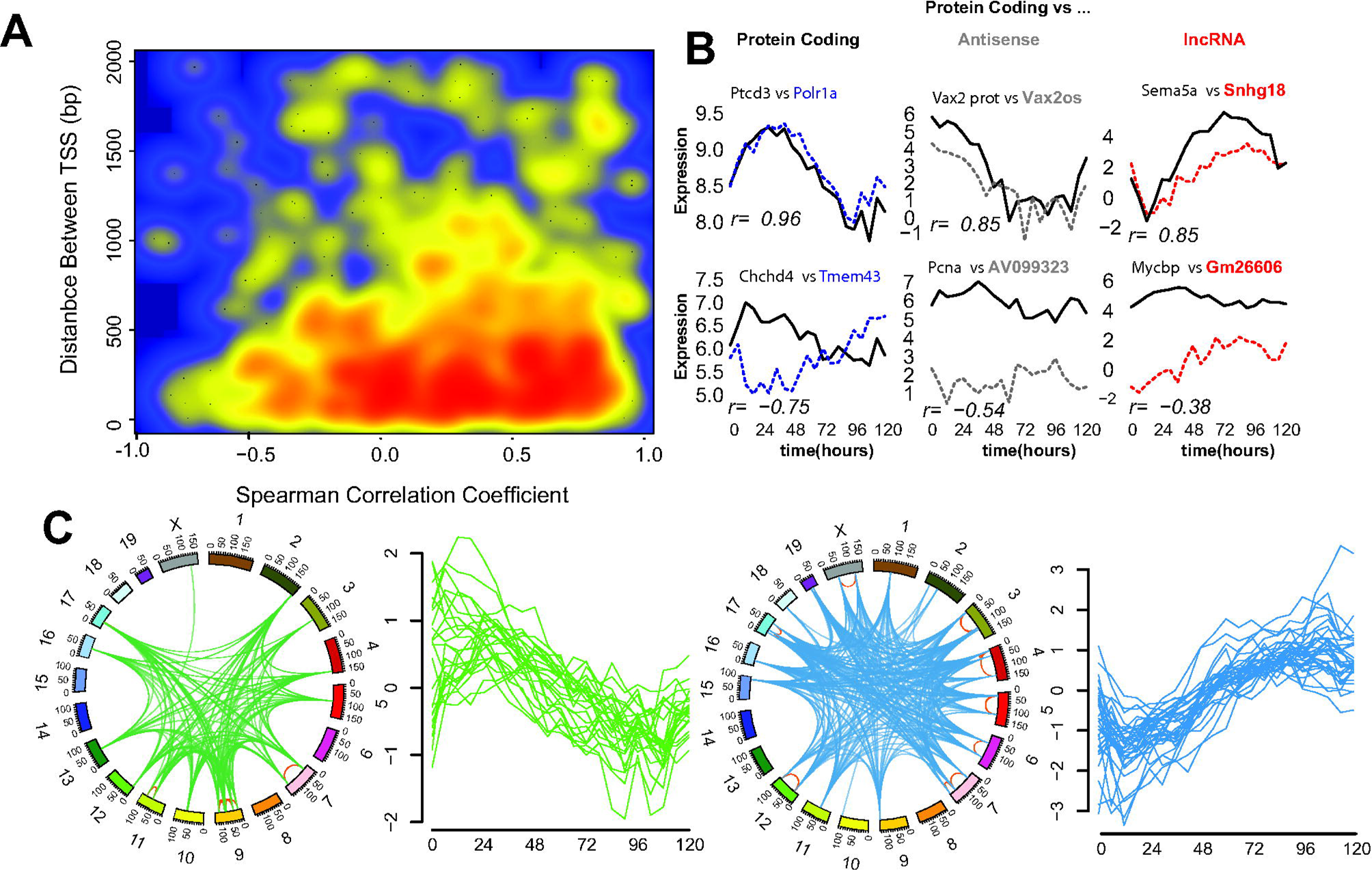
Analysis of gene coexpression patterns using augmented temporal resolution. (**A)** Smoothed scatter plot showing the correlation coefficient across the time course vs. distance between transcriptional start sites (TSS) of bidirectional gene pairs. Blue indicates no gene pairs; yellow and red indicate increasing numbers of pairs sharing similar properties. (**B)** Expression patterns of example bidirectional genes of the same or different gene biotype. Spearman’s correlation coefficient is reported for each pair. (**C)** Genomic location (circos) and expression pattern (line plot) of two independent co-expressed groups of 5 or more contiguous genes sharing correlated expression (r>0.5).

To investigate whether the strong correlative potential between gene pairs could facilitate the identification of regions of the genome that are coordinately regulated (61), we scanned across the genome for regions containing five or more contiguous genes that were coexpressed (r>0.5). This revealed 59 regions with a mean size of 821 kb-each containing 5-14 genes (mean of 6) genes. The majority of these regions were each contained within a single topological associated domain(62) (Supplementary Figure S4F), increasing the propensity for a common regulatory architecture. Evaluation of gene-expression patterns across these clusters revealed evidence of high co-expression at both the inter- and intra-chromosomal levels (Supplementary Figure S4G). We assembled a map of regions of the mouse genome displaying high levels of clustered co-expression (Figure 3C) by comparing the expression profiles of the regions. Two independent modules were identified with distinct decreasing (green)- and increasing (blue) expression patterns with differentiation. Given the independent location and expression patterns of these clusters, we suggest these regions may form core expression-factories of cellular differentiation. In support of this notion, this analysis identified the gene cluster-associated with the “increasing module”-containing the imprinting locus of H19, Igf2, Tnn3 and Mrpl23(63) (Supplementary Figure S4H); previously shown to be activated in concert during early stem cell differentiation(64).

These investigations illustrate how analysis of high-resolution temporal transcriptomic data provides an independent and convenient approach (relying only RNA-Seq) to guide the partitioning of the genome into regulatory domains.

### Increased temporal resolution refines the noncoding landscape of mESC differentiation

Having shown that rapid changes in lncRNAs are a key feature of ES differentiation, and that co-expression analysis is a powerful tool for understanding gene regulation with augmented temporal resolution, we sought to unravel the roles that lncRNAs might play in ESC differentiation.

Analysis of gene annotations yielded confident expression data for 588 lncRNA genes at six-hourly resolution (520 for 24-hourly, Supplementary Table S1). Indeed, added temporal resolution increased information of all noncoding transcript biotypes indicating that a proportion of these genes were only present for a short duration in this system. Clustering lncRNA expression patterns with time-dependent protein coding gene expression showed that lncRNAs were enriched at lower expression levels and shared related expression profiles to protein coding genes (Figure 4A). This relationship was further examined whereby K-means clustering of these expression profiles compared to clustering of a similar number of time-dependent protein coding genes (Figure 4B, Supplementary Figure S6A) revealed clusters of lncRNA genes resembling gene expression patterns associated with stemness (cluster a) primitive streak formation (cluster b) and WNT signaling (cluster c)(14). Determining the role that these lncRNAs play in these processes will be important in understanding the molecular events underlying cell differentiation.

**Figure 4.**
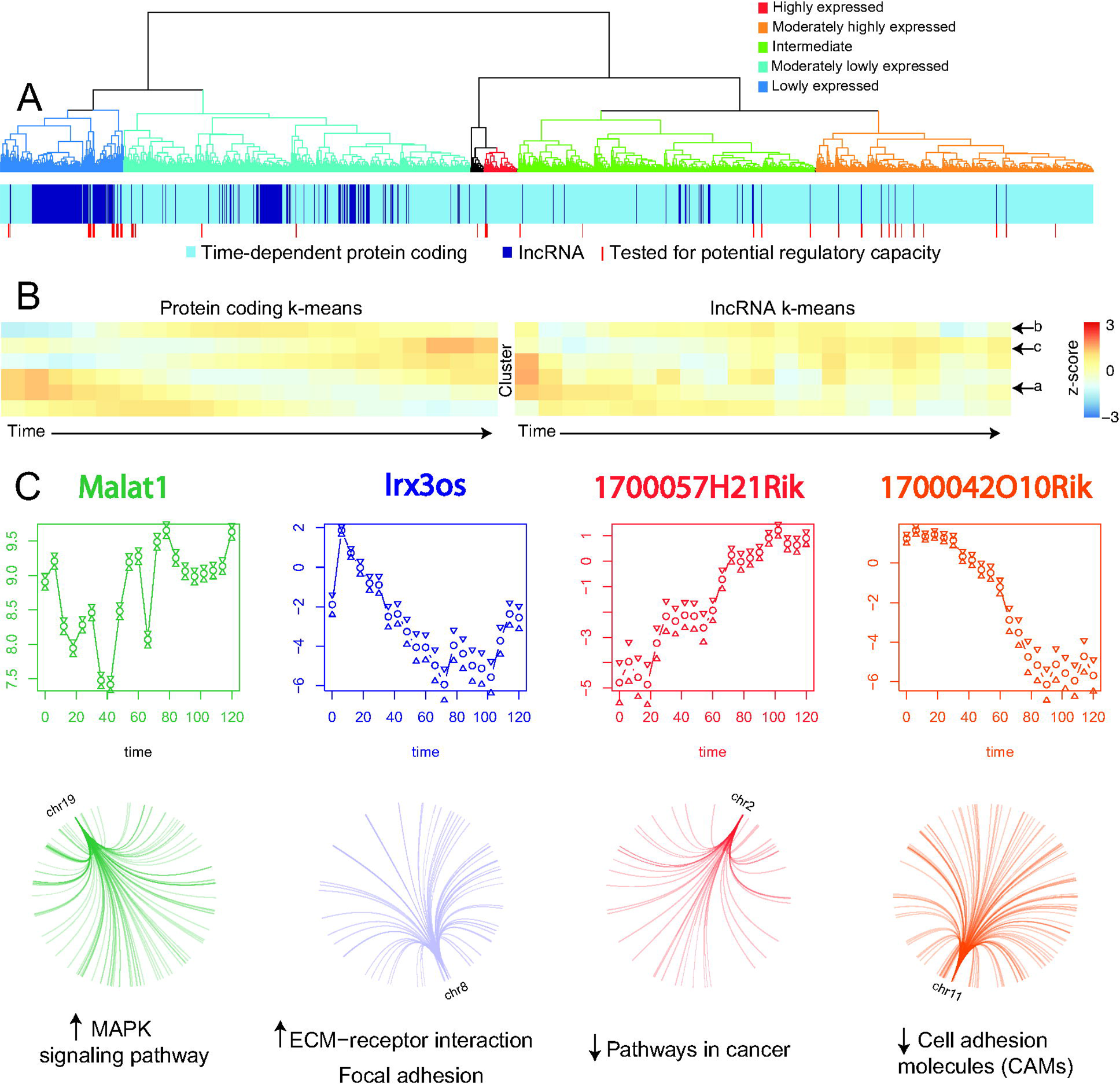
Augmented temporal resolution of ncRNA expression in cellular differentiation. **(A)** Hierarchical clustering of lncRNAs (dark blue) with time-dependent protein coding genes (light blue) by their expression patterns over time. Dendrogram was manually colored to reflect gene expression levels of the top-level clusters. (**B)** K-means clustered expression profiles of protein coding genes compared to the same number of lncRNA gene expression clusters. Common profiles are marked with arrows. (**C)** Expression profiles of four lncRNAs predicted to have regulatory roles in ES development as well as the genome location & pathways enriched in their gene targets. Malat1 and IRX3os display a positive association with their targets, whereas 1700057H21Rik and 1700042O10Rik have a putative repressive impact.

As lncRNAs often exert their function through guiding or assembling transcriptional machinery, we sought to identify potential regulatory lncRNAs in this system. We selected 50 highly or variably expressed lncRNAs (Figure 4A) and tested for evidence of gene regulatory behavior across the transcriptome. Since lncRNAs typically exert their function as a transcript, we set a maximum time offset of 18 hours to avoid secondary (altered protein level) effects and examined patterns in the predicted gene targets of lncRNAs (r>0.8, p<0.05, divided by positive or negative associations). Reactome pathway analysis revealed that 11 of these lncRNAs (including well characterized lncRNAs, Supplementary Figure S6B&C) were potentially involved in regulating networks of genes associated with key developmental processes (p.adj<0.05, Supplementary Figure S6C). These analyses assigned target gene networks consistent with characterized lncRNA biological functions for Malat1 (oncogenic(65)), Neat1 & Rian (association with gene repression(66)) and Meg3 (tumour suppressor(67)). Interestingly, these data suggest that the pro-tumorigenic function of Malat1 may be mediated through facilitating the increase of MAPK signaling molecules. Importantly, these data also provide testable evidence for seven previously uncharacterized lncRNAs role in ES development and describes a map of regulatory interactions driven by lncRNAs (Figure 4C) whereby lncRNAs can affect gene expression across the genome. The identification of lncRNAs with a predicted biological role is important for unraveling lncRNA function, providing candidate functional lncRNA and providing a level of molecular detail that is currently lacking in many lncRNA studies.

## Discussion

Transcriptional regulation of key biological events is a key feature in understanding the complexity of cellular processes. Here we describe a detailed transcriptomic resource for research in cellular development, a framework for unraveling this detail and identifying new targets for analysis. We also present a comprehensively detailed survey of noncoding transcripts throughout early stem cell development. We have identified many previously uncharacterized noncoding RNAs with potentially pivotal roles in cellular differentiation. This will provide a valuable tool for researchers unraveling the transcriptional complexity of cellular differentiation.

### Increased interpretive power

The understanding of molecular events underlying the departure from pluripotency has been determined by the extant knowledge of how biological functions are exerted – often measured at 24 hourly or greater intervals. We hypothesized that interpretations of this model were missing detail in light of evidence indicating the unforeseen dynamics in RNA biology and regulation. By probing this detail with finer time distinctions, we show that gene expression profiles of well-characterized genes display significant variation of expression levels and that such variations are manifest in a significantly more complex gene regulatory framework. This is consistent with a reduction in temporal aggregation bias (18) and highlights early array-based investigations in yeast demonstrating the importance of sufficient temporal resolution in understanding gene expression patterns (68). As such, much detail is likely missing from other systems that involve a change in phenotype or cellular behavior. With large-scale transcriptomic analyses becoming increasingly accessible, it is opportune to revisit other well-studied transitions with the view of improving understanding and applicability of their results rather than relying on presuppositions about gene expression patterns (69).

### Insights into short bursts of transcription

We have shown the benefit of frequent sampling over time in observing the transcription of genes that are observable only within sub-24 hourly windows. This approach highlights the importance of taking into account the presence of short-lived transcripts and shows that cells express more of the transcriptome in a time-dependent fashion. To this end, we have identified rapid changing and periodically expressed genes, which we term short-lived (slRNA) and cycling (cycRNA), that were unobservable outside this framework. That many slRNAs exhibited changes in expression over the first 24 hours of differentiation is consistent with rapid initial cellular response to stimuli (20,70). Indeed, it is likely that significant gene expression changes-especially noncoding-occur on timeframes shorter than those presented that may not be amenable to optimal timepoint prediction strategies (69). By probing deeper into time-dependent gene transcription-possibly by interpolating available datasets-(68) it will be possible to uncover further complexity underlying cellular plasticity and gene regulation. These observations reinforce the concept that adequate temporal resolution is vital for describing biological transitions-for example in dissecting primary from follow on effects in gene knockdown studies – and that end-point analysis likely does not reflect the complex biology of phenotype changes.

### Insight genome organization and regulation

Similarly, by using time to separate the order of gene transcription, we have been able to predict local gene regulation across the genome. We have been able to observe concerted gene expression (in trans) of hundreds of genes separated by large genome differences (in cis). Typical studies of this nature involve correlative analysis requiring large samples sizes and resources (71). We have instead leveraged the time axis to achieve these as well as discriminate driver from passenger molecular events. This has allowed the estimation of the time delay for changes in expression of regulatory molecules to manifest in changes in their target gene transcription and we have been able to unravel a potentially complex network of gene profiles responding to lncRNA transcription. Finally, we have been able to use an integrated biological system to draw strong associations in trans relationships with bidirectional promoters. Typically these associations are observed by using thousands of gene expression profiles, yet here we have been able to do so with only 42.

### General experimental considerations

The design and interpretation of time course experiments has been of great interest over the past decade (18,69) and they have been used effectively to elucidate transcriptome expression and regulation in many organisms (21-24). Furthermore, improvements in sequencing technologies are making the dynamics of larger and complex genomes more available to closer inspection. By probing transcriptional complexity in mouse ES development, we have gained insight into many areas of molecular inquiry. Using uniform dense sampling enables strong gene expression relationships to be drawn whilst simultaneously facilitating the dissection of expression ordering and kinetics. Importantly these data show that substantial changes in gene expression cannot be inferred from coarse time-points and that the continuous representation of gene expression data in many developmental time courses obscures detail. Therefore the assumptions made when choosing time points for these kinds of studies (such as how long a biologically significant event takes to occur) need to be re-evaluated; RNA and protein turnover is extremely rapid (72) and transcriptional responses are extremely rapid (20,68,70) and can be transient (73). Our data suggest that dense profiling will yield more insights into reprogramming and that a real-time picture is yet to be achieved (Supplementary Figure S7A). Furthermore, using temporal approaches to augment single cell transcriptome studies such as dissecting cellular heterogeneity (31) and lncRNA expression patterns (74) similar to the method employed in (19) may allow the temporal tracking of single cell alterations over time.

Analysis of high-resolution temporal transcriptomic data reveals an unprecedented level of regulatory complexity and presents a tantalizing opportunity to revisit and bring new insight into other clinically or biotechnologically significant biological transitions. In designing these experiments it is important to choose the approach to match the aim. For example, gene knockdown experiments using siRNAs may benefit from early time point transcriptomes for dissecting primary from secondary or tertiary effects. Uniform temporal sampling simplifies the interpretation of temporal correlations in gene expression whereas focus on early responses will necessarily require rapid initial time point selection with tracking samples. The frequency of collection will necessarily depend on the duration of the response and practical and financial considerations. Increasing density will necessarily increase the correlative power of the study without negatively affecting the observation of transiently expressed genes (Supplementary Figure S7B & (75)). However, replication ensures uniformity of the biology underlying the process in question (figure 1B), thus enabling confident dissection of transient or periodic gene expression patterns from noise.

## Acknowledgements

The authors acknowledge Kenneth Sabir and Ruth Pidsley for reviewing the manuscript; the Garvan Foundation and the Peter Wills Bioinformatics Facility for providing facilities and Agilent Technologies for providing RNAseq kits. BG acknowledges constructive feedback from Aaron Statham, Mark Pinese, Nenad Bartonicek, Jesper Maag and Quek Xiucheng

## Abbreviations

lncRNA: logn noncoding RNA
MPS: massively parallel sequencing
ESC: embryonic stem cell
PCA: principle components analysis
TF: transcription factor
slRNA: short-lived RNA
DEX: differential exon
cycRNA: cycling RNA

## Declarations

### Ethics approval and consent to participate

Not applicable

## Consent for publication

Not applicable

## Availability of data and material

Data has been deposited into GEO: The Reviewer access link is: http://www.ncbi.nlm.nih.gov/geo/query/acc.cgi?token=cnglummejbkltyj@acc=GSE75028

## Authors’ contributions

BG wrote the manuscript, assisted study conception, performed the analyses and library preparations (assisted by DK). MD conceived the study and assisted writing the manuscript. BS performed some analyses (de-novo assembly and PC deconvolution), designed the web portal, assisted with figure composition and reviewed the manuscript. SC, FG and DK performed lab work and reviewed the manuscript. AP provided biological samples and facilities

## Authors’ information (optional)

Correspondence and requests for materials should be addressed to

## Supplementary Figures and Legends

Supplementary Table S1: Counts of Genes by biotype

Supplementary Table S2: Periodic Genes

**Supplementary Figure S1.**
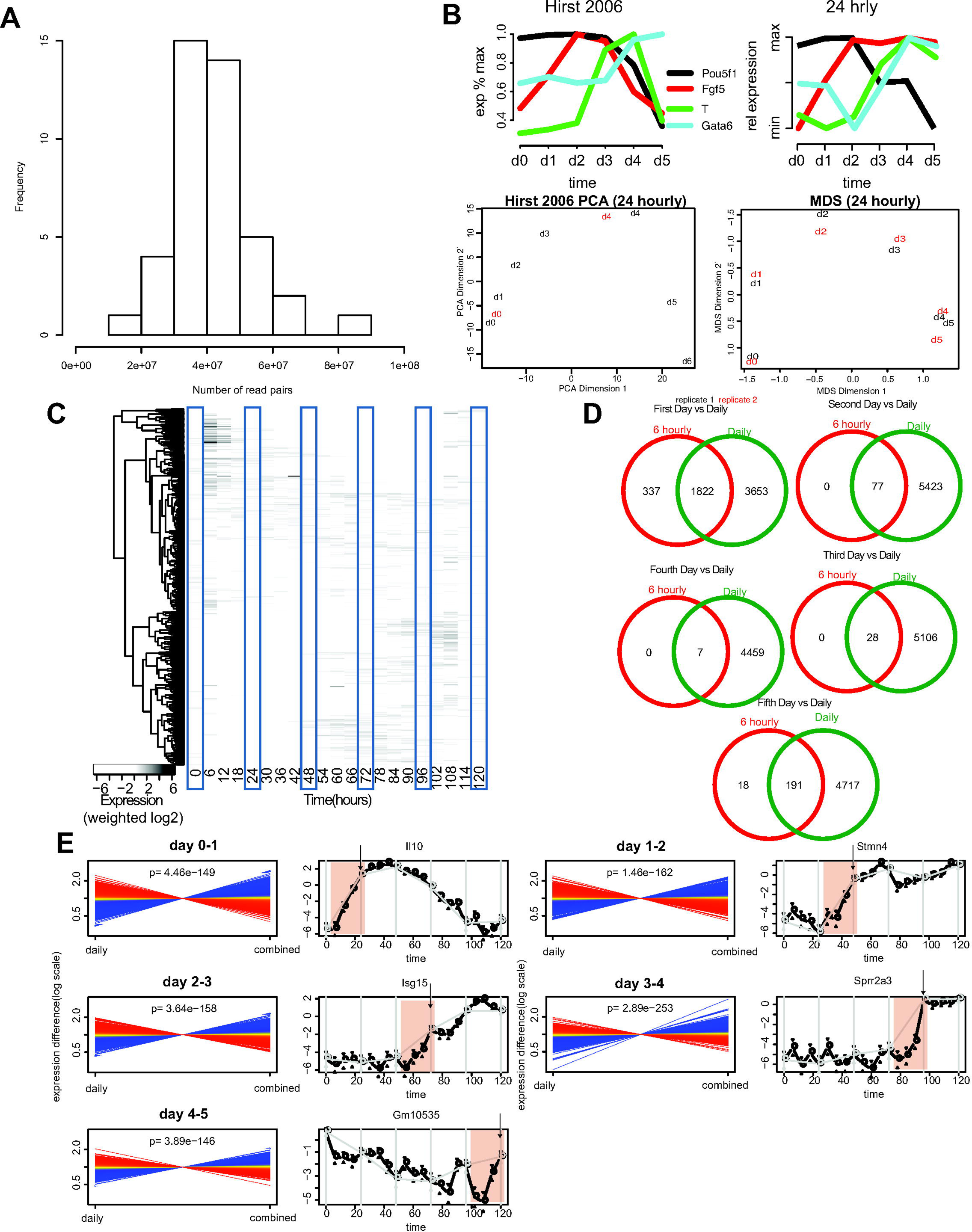
Global evaluation of high-resolution transcriptomic data. (**A)**Histogram of mapped read number distribution per sample (pooled from biological replicates). (**B)** Comparison of expression levels and principle components analysis measurd 24 hourly between this study and Hirst 2006 **(C)** Heatmap of expression levels for genes only expressed outside of 24 hourly timepoints, clustered by expression pattern. **(D)** Evidence of differential expression within one 24 hour period vs. any change across all 24hourly times (p <0.0001). (**E)** Comparing whether the 24 hourly measures “summarize” that 24 hour window by comparing mean expression for that window with the 24 hour time point

**Supplementary Figure S2.**
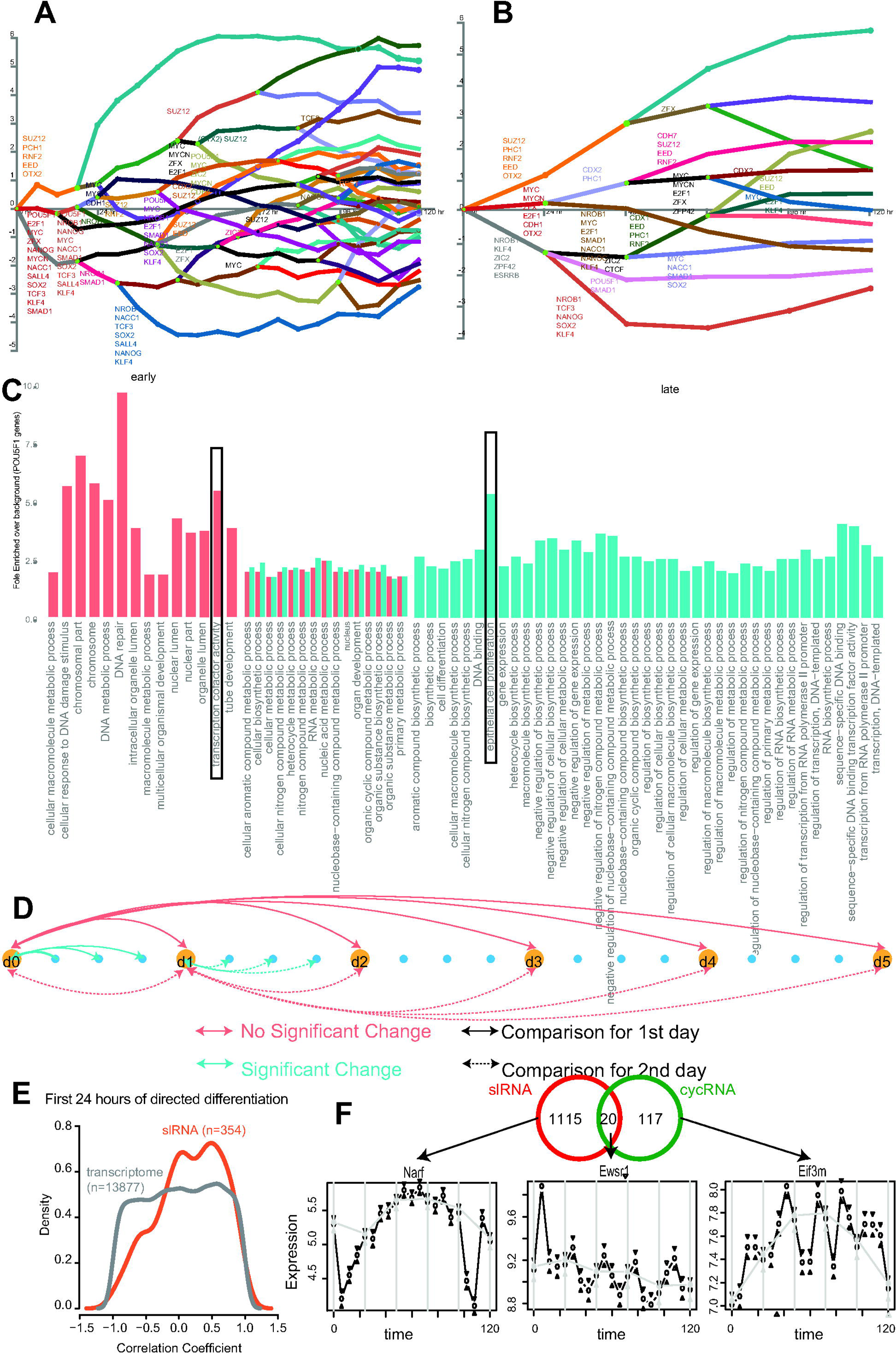
Highlighting unique knowledge gained from increased temporal resolution. **(A &B)**Fully annotated DREM schematic of estimated TF activity of key ESC related TFs at 6hourly (A) vs. 24 hourly (B). (**C)** GO term enrichment (adjusted p<0.05) for genes corresponding branch points designated as early (co-observed with change in POU expression) and late (observed after POU5f1 Expression changes) highlighted by the blue boxes in Figure 2A. Black boxes represent similar terms identified in figure 2D. (**D)** Schematic of differential expression analysis design used to identify slRNAs. (**E)** Correlation of slRNA expression in (De Kumar et al. 2015). **(F)** Comparison of slRNAs and cycRNAs. Venn diagram of the overlap observed and examples from each class.

**Supplementary Figure S3.**
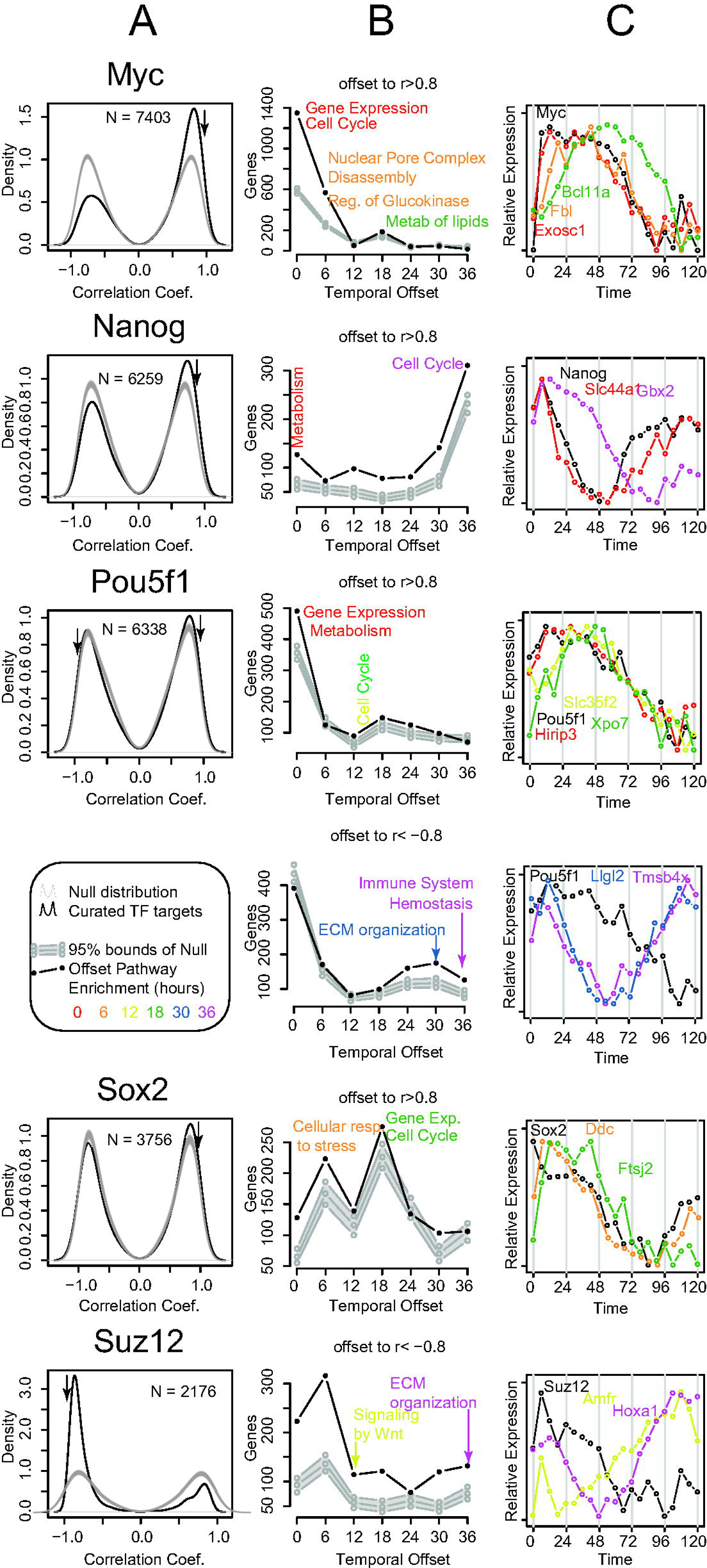
Temporal offsets in transcription factor (TF)-target gene expression. (**A)** Curated TF/gene targets were downloaded from chea (http://amp.pharm.mssm.edu/lib/chea.jsp) for Myc, Nanog, Pou5f1, Sox2 and Suz12. Expression of target genes were tested for correlation with their TF at different temporal offsets (0-36 hours) and compared to 500 random selections of the same number of genes (Null). Where absolute correlations of predicted targets exceeded the null distribution (arrow), (**B)** the number of genes achieving a maximal absolute correlation of >0.8 and the offset required to reach these maxima was plotted against the 5th and 95thquantiles of the same results from the null distribution. Where the number of target genes exceeded the null distribution, the lists of genes in each offset were tested for enrichment of Reactome pathways relative to the total predicted target list (enrichment). (**C)** Example expression patterns of genes displaying these attributes were plotted.

**Supplementary Figure S4.**
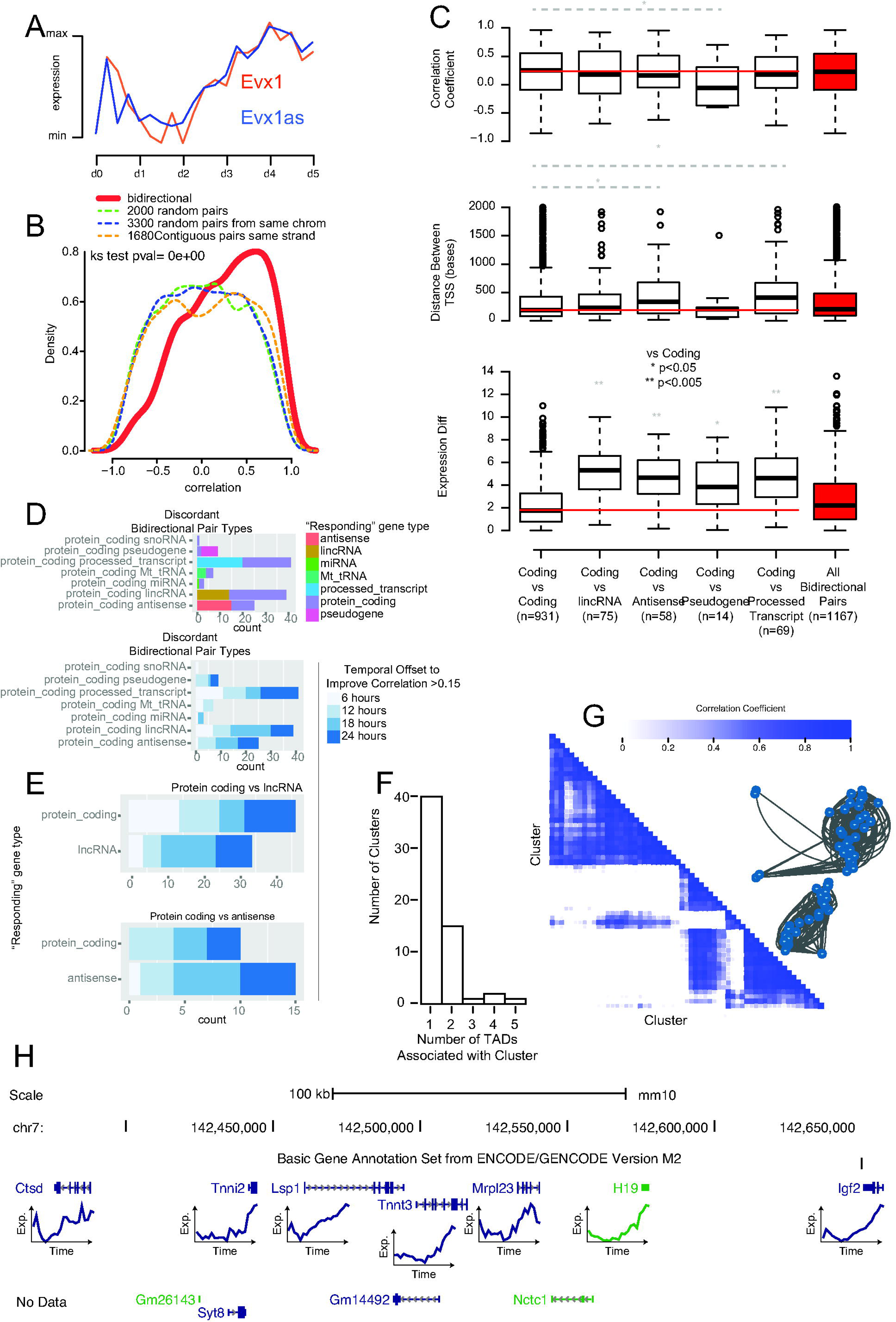
Bidirectional and co-expression analysis of mouse ES development. **(A)**Expression profile of EVX1 and its antisense (and positively correlated) transcript EVX1AS- the peak at 6-18 hours has not been observed previously. (**B)** Distribution of correlation coefficients of bidirectional gene pairs (red) compared to similar numbers of randomly chosen genes pairs, randomly chosen genes from the same chromosome and, randomly selected neighbouring genes (dotted lines). ks=Kolmogorov–Smirnov test (bidirectional vs. random neighbouring gene pairs). (**C)** Characteristics of bidirectional gene pairs (Correlation coefficient, Distance between TSS and Difference of median expression (log scale) based on annotated gene-biotype. (**D)** Counts of bidirectional gene pairs of differing biotypes achieving an improved correlation coefficient of >0.15 (to at lease 0.25) over that at time zero, colored by the biotype of the “following” gene or by the temporal offset required to achieve the improvement. (**E)** Comparison of responding gene biotype to the temporal offset for lincRNA and antisense biotypes. (**F)** Number of topological associated domains (TADS, HindIII data mapped to mm10 using liftOver from mm9) associated with each co-expressed gene cluster. (**G)** Clustering of co-regulated gene clusters by correlation coefficient visualized by network diagram and hierarchical clustering of the correlation matrix. (**H)** The imprinted H19/IGF2 cluster identified as a co-expressed gene cluster with gene expression data for measured genes. Some genes did not have expression data (no data).

**Supplementary Figure S5.**
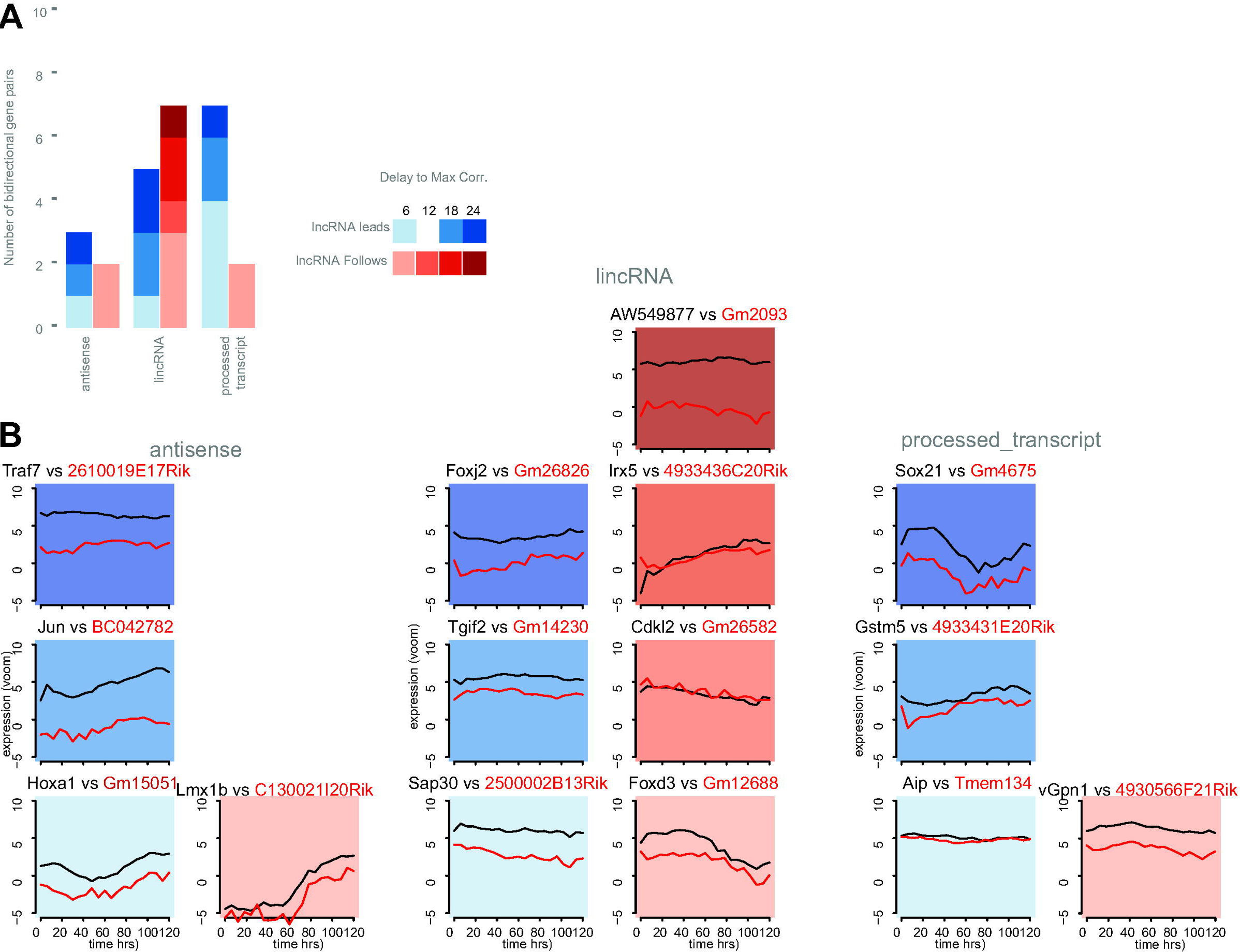
Temporal relationships of highly correlated coding-noncoding bidirectional pairs. **(A)**Bar chart of the temporal offset required to reach a maximum correlation >0.8 and whether the noncoding gene preceded the protein coding gene or vice versa. (**B)** Example gene expression profiles of bidirectional paired gene over the time course. Gene profiles are arranged and colored as the bar chart.

**Supplementary Figure S6.**
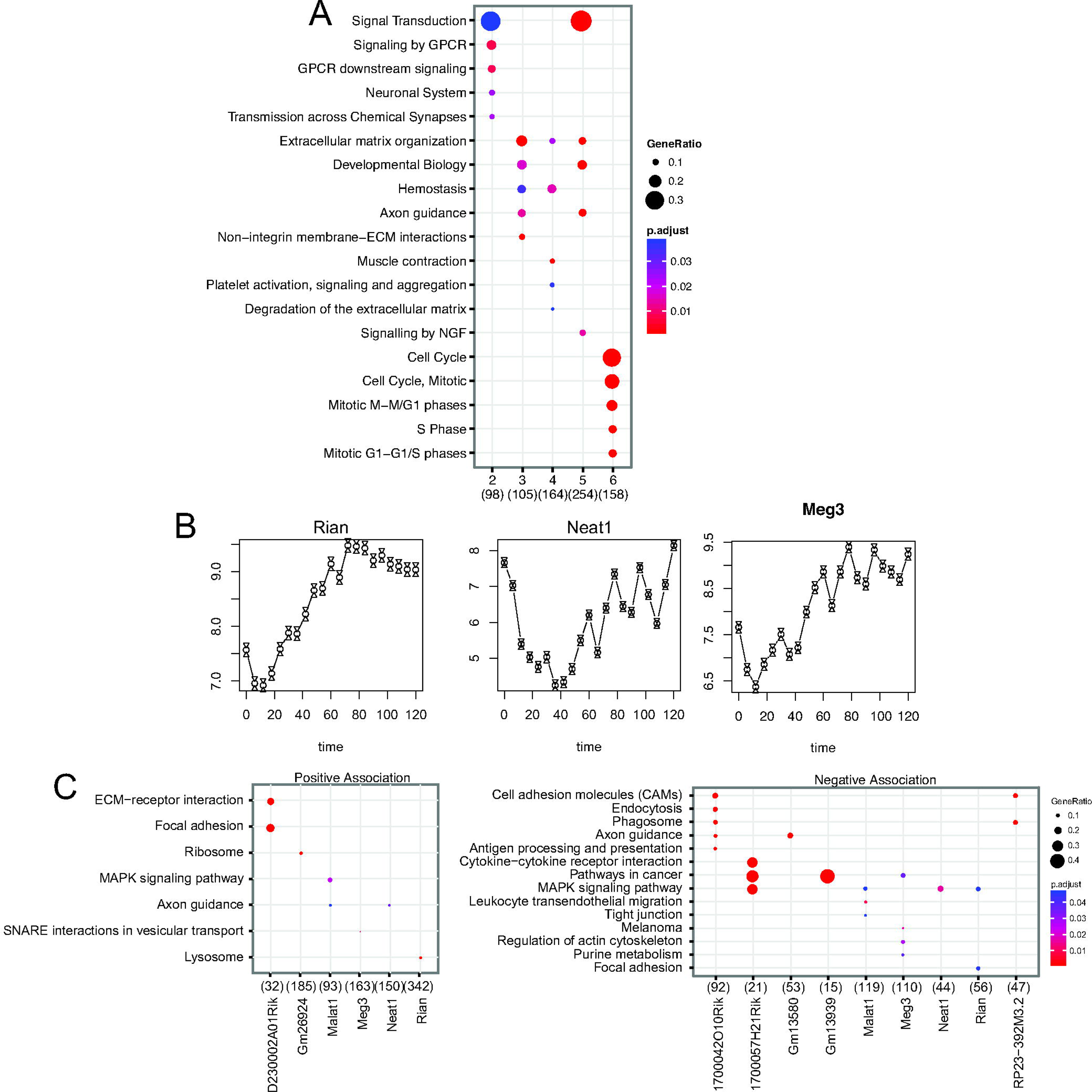
LncRNAs and their role in ES development. **(A)**Reactome pathway enrichment for 5/6 k-means clusters of time-dependent protein coding genes. (**B)** Expression profiles for characterized lncRNAs described in text. (**C)** Reactome pathway enrichment for putative gene targets positively or negatively associated with candidate lncRNAs (top 4 pathways, enrichment adj. pval.<0.05)

**Supplementary Figure S7.**
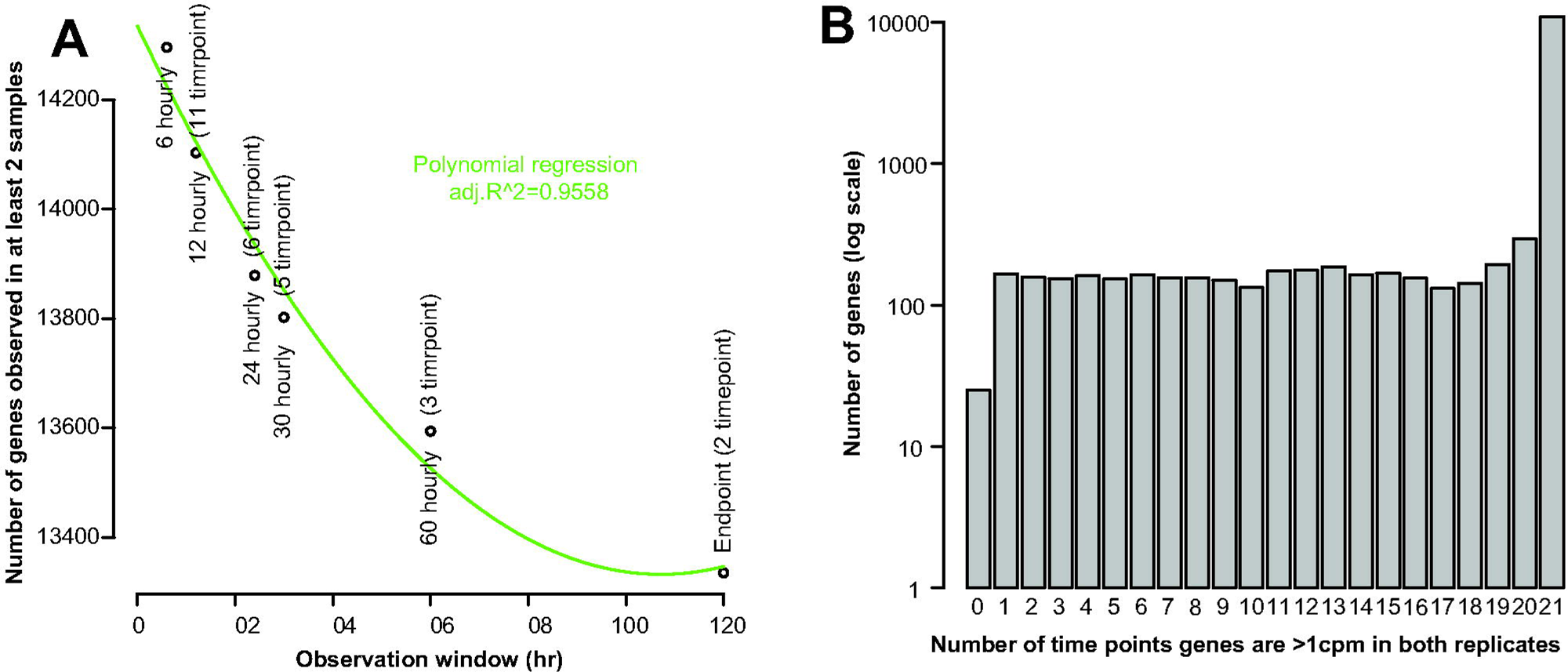
Sampling density impact on gene expression observations. **(A)**The impact of increasing temporal resolution on the number of genes observed to be expressed. **(B)** The number of conditions in which each gene observed is expressed above background in both replicates across the time course. ~150 new genes are observed at a single timepoint.

